# Resolving Genome-to-Phenotype Links in Bacteria: Machine-Learned Inference from Downsampled k-mer Representations

**DOI:** 10.64898/2026.02.18.705352

**Authors:** T. Regueira, C. Quaglia, O. Lund

## Abstract

Standard approaches to bacterial phenotyping often treat the entire genome as the fundamental unit of information, resulting in high-dimensional inputs that may contain significant redundancy. Consequently, current bacterial phenotyping techniques typically rely on the assumption that entire sequences are required for accurate predictions. While downsampling based on min-hashing or prefix filtering has been used for clustering, its utility as a direct input for predictive machine learning remains underexplored. Here, we show that a novel prefix-based downsampling algorithm can reduce the size of genomes while maintaining relatively high predictive accuracy on phenotype prediction tasks. By combining a prefix reduction strategy with the specificity of short k-mers, we developed a method to downsample entire genomes into k-mer frequency matrices and *k-mer-on-a-string* representations. We found that ensemble models, such as Random Forest and Gradient Boosting, trained on k-mer frequency matrices from downsampled genome representations outperformed more complex deep learning architectures with the same downsampled representation, particularly on datasets with limited data or highly similar genomes. We were able demonstrate explainability by tracing back the k-mers with the most impact on the models to genes coding for the specific phenotype. Our results demonstrate that downsampling genomic data can yield models with good predictive power thus establishing an alternative when using full genomes is infeasible. We present an approach that offers relatively high performance on bacterial phenotyping tasks and demonstrates a path forward towards lightweight Genome Language Models that will enable analysis of entire genomes.

## 1 Introduction

Bacteria are incredibly diverse, live in almost any geography, and are extremely good at adapting to new environments (Fenchel, 2003; Lai & Cooper, 2021; Byrne et al., 2025). It has been estimated that more than ∼ 10^12^ microbial species exist (Locey & Lennon, 2016). When bacteria adapt to new environments, these adaptations are often the result of changes in their genomes. These changes are distributed across the genome and on different genomic elements. Functional genes are contained in operons, whereas resistance and virulence genes most often are located on genetic elements such as plasmids and acquired through horizontal gene transfer (Koonin, 2009; Rodríguez-Beltrán et al., 2021). Additionally, it has been found that bacterial gene families usually are found in specific locations on the chromosomes (Hu et al., 2025). This means that the structure of the bacterial genome and genomic elements can help us interpret and infer bacterial phenotypes.

Prediction of phenotypes from bacterial genomes is a challenging task, made harder by a sparsity of sequenced bacterial genomes with phenotypic annotations. A diverse set of species is needed for phenotype prediction tasks, to create Machine Learning models that are able to explain the variance in the genomes, and be able to generalize to species not seen in the given set of genomes.

To enable phenotype prediction from genomes, the genomes must be encoded into a representation that the Machine Learning model understands. One approach for encoding DNA sequences as inputs interpretable for a Machine Learning model is the fixed-length bag-of-k-mers approach, where overlapping k-mers are sampled using a sliding window over the genome, and every k-mer is counted in a k-mer absence-presence or counts matrix, possibly normalized. This approach has been used with success to predict regulatory sequences (Ghandi et al., 2014) and to uncover the regulatory architecture of maize (Mejía-Guerra & Buckler, 2019). Representing an entire bacterial genome as a frequency matrix is a relatively crude representation of the genome, but it is manageable for many models, like Random Forest and Gradient Boosting. Up to a K-mer length of around 10 all possible K-mers can have their frequency or presence/absence encoded as input while keeping the number of inputs below 1 million. For longer K-mers it may only be feasible to encode a one hot encoding for each unique K-mer seen in the training set, rather than all possible k-mers of that length. Representations of entire genomes at a single nucleotide scale require a longer context length than what state of the art transformer architectures are capable of, since many bacterial genomes are more than 5 million base pairs (Mbp) long, while the context length of Transformer models typically is around 12 thousand base pairs (kbp) (Dalla-Torre et al., 2025). However, a genomic representation for predicting phenotypes does not need to be a full DNA sequence. Instead, several studies have shown that a genome can be divided into proteins and each protein encoded as, e.g., a one-hot vector or an embedding. Then the collection of proteins is used to model the genome and predict bacterial phenotypes with high accuracy (Dalla-Torre et al., 2025; Wiatrak et al., 2025). A recent approach is to use the newly designed Mamba (Dao & Gu, 2024) or Hyena DNA (Nguyen et al., 2023) machine learning architectures, which can model much longer sequences (+1 Mbp) with a performance close to that of the traditional transformer architecture. However, the training of any of these genome language models is computationally expensive to run and is still not able to encode entire genomes. Here we will examine whether it is possible to downsample DNA sequences in such a way that their structure and information content is mostly preserved while the overall data size are dramatically reduced, thereby reducing computations needed when modeling the genomes.

MinHashing is a common approach to analyzing genomes and estimating genome distances. MinHashes were originally used to compare similarity between the content of web pages (Broder, 1998). Similarly, a genome can be represented as a list of overlapping k-mers, and each k-mer in the genome can be represented as a hash using a hash function. The MinHash function decreases the number of k-mers by selecting k-mers with the numerically N smallest hashes. This significantly decreases the amount of sequence information used for the comparisons while still being able to compare sequence similarity, clustering sequences, and classify phenotypic traits of genomes. Mash (Ondov et al., 2019) and SourMash (Irber et al., 2024) are widely used tools based on MinHashing and used for this very purpose in bioinformatics, where they enable clustering of large sequence databases (Ondov et al., 2019). Downsampling with the use of a min-hashing algorithm results in a massive compression factor compared to the raw FASTA files (Ondov et al., 2019).

We have previously developed a “prefix downsampling algorithm”, which has similarities to the MinHashing algorithms, but also some key differences(Larsen et al., 2014). We will here investigate the use of down sampled genomes using the prefix strategy as input for machine learning methods. In Larsen et al. (Larsen et al., 2014) the first five nucleotides were fixed, and only the k-mers matching the first five nucleotides were added to the database containing 16-mer nucleotide sequences. This resulted in a approximately factor 1000 downsampling of k-mer space and enabled the prediction of bacterial taxonomy.

The prefix downsampling algorithm enables the compression of a large bacterial genome to a much smaller representation while retaining gene order and can be seen as “lossy compression”, similar to the SourMash algorithm (Irber et al., 2024). The degree of down-sampling is controlled by the prefix length, k, which determines the specificity, and the suffix length, l, which controls the amount of information we keep per match and thus the amount of total DNA sequence saved to the downsampled representation.

After compressing the sequences to the downsampled representation, they can be converted into a numerical representation for the Machine Learning models. A common, simple representation of a genome is to create a binary presence/absence matrix of the k-mers / tokens, which can easily be fed to many different types of Machine Learning models. A similar approach to this has been used with success to predict antimicrobial resistance (Aytan-Aktug et al., 2020). Alternatively, the k-mers can be represented in a counts or frequency matrix. A different approach would be to feed the k-mers to a machine learning model as *k-mers-on-a-string*, where each k-mer is represented as, e.g., a one-hot encoded vector or integers. This genome representation keeps the order of the down-sampled k-mers, which we argue will benefit some models. This type of representation can be used in Convolutional Neural Networks (CNNs), Recurrent Neural Networks (RNNs), and in the Transformer architecture.

Here we will investigate whether down-sampled genome representations can be used to create high-performance machine learning models. We will combine prefix downsampling strategy from (Larsen et al., 2014) with the short k-mer approach from (Aytan-Aktug et al., 2020) and compare the k-mer frequency matrix to a k-mer-on-a-string approach in a Machine Learning context. Additionally, we will evaluate how well different machine learning architectures predict various bacterial phenotypic traits.

## 2 Methods and Materials

### 2.1 Data

Two different dataset sources were used to develop and benchmark bacterial genome representations. The first dataset is used to train and evaluate the prediction of bacterial phenotypic traits from the *Bacformer* paper (Wiatrak et al., 2025). The second dataset is used for predicting resistance towards the antibiotic gentamicin in *Escherichia coli*.

#### Dataset 1: Prediction of bacterial phenotypic traits

In the bacterial phenotypic traits study, we use a dataset originally compiled and used in the *Bacformer* paper (Wiatrak et al., 2025) to design and train different Machine Learning models. The dataset consists of 24,462 bacterial genomes extracted from Genbank, containing 15,477 species across a diverse set of phenotypic traits as labels. Each row is a genome encoded as DNA with spaces separating the contigs. This dataset is a collection of 50 parquet files, each containing around 500 genomes, and a separate metadata file containing the labels for the different phenotype prediction tasks.

#### Dataset 2: Prediction of gentamicin resistance in *Escherichia coli*

The gentamicin resistance dataset was obtained from the BV-BRC database (Olson et al., 2023), consisting of 966 different randomly selected *E. coli* genomes, of which 423 are classified as gentamicin resistant and 543 as susceptible to gentamicin, making the dataset relatively balanced. The dataset consists of individual fasta files for each genome and a separate metadata file with the label resistant or susceptible.

### 2.2 Data preparation and downsampling

To prepare genomes as inputs for training the machine learning models, the dataset files (.fasta or .parquet format) are loaded, and genomes are treated one-by-one using the “prefix downsampling” approach, where a short DNA-string, the “prefix”, is selected and is slid across the genome, looking for matches between the short prefix and the genome. When a match is found, a piece of DNA of size n after the match, the “suffix”, is appended to a list of suffixes, which is then saved as the downsampled genome fig. 1. The size of the prefix and suffix can be used to control the degree of downsampling.

**Figure 1:**
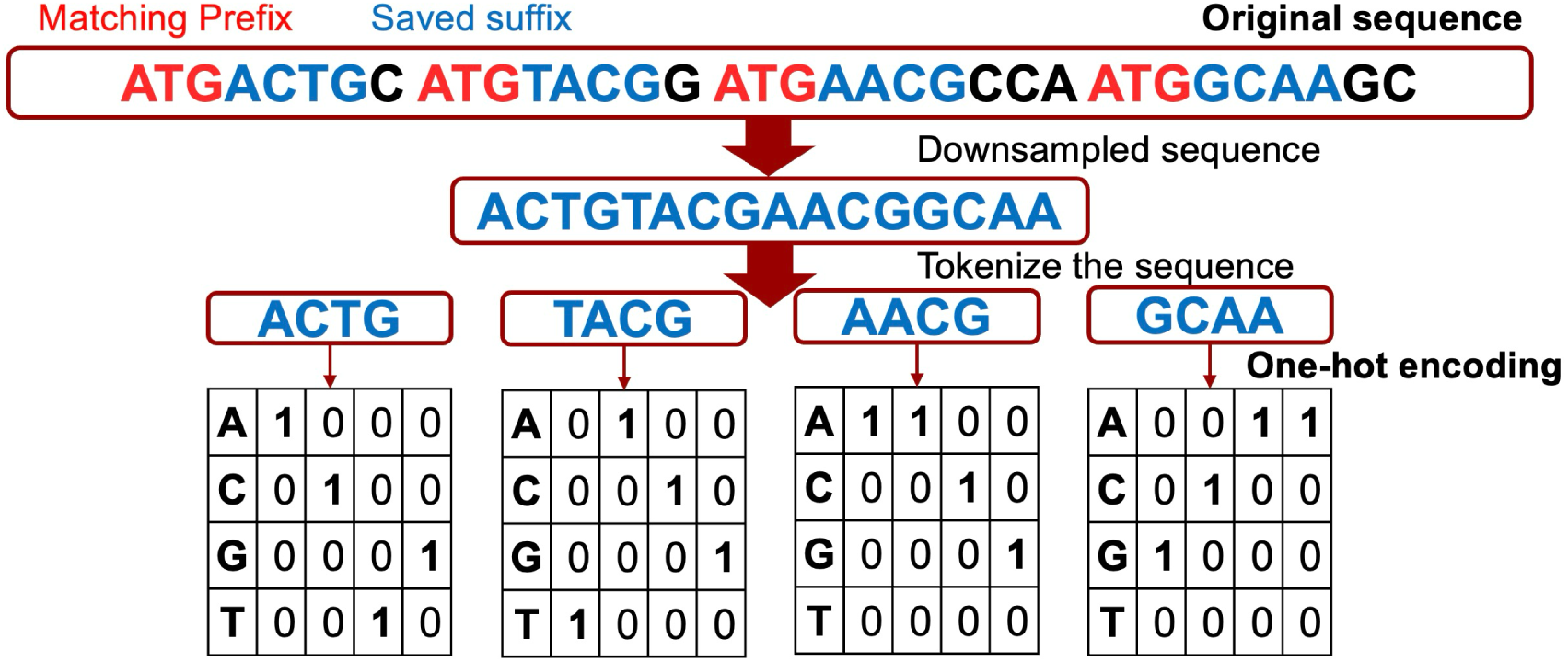
Workflow for *K-mers-on-a-string* approach. The piece of DNA sequence that follows the matching prefix is saved to the downsampled sequence. In the one-hot encoded approach, a sequence is tokenized and encoded using the one-hot encoding approach, either using tokens of 1 or more nucleotides. The ESM-C approach translates the entire DNA sequence to amino acids, including stop codons, and encodes each amino acid as a vector. To obtain the full genome representation, the mean of each row in the vectors is computed, resulting in one vector per genome.

### 2.3 Genomic encodings

After downsampling, the genomes are processed in one of two ways, depending on whether the data is used as input for the ensemble models or for the more complex machine learning models.

For the Random Forest and HistGradientBoosting models, a simple k-mer counts or frequency matrix is used to encode the genomes by counting the occurrence of each DNA suffix and dividing by the total number of matching suffixes per genome (table 1).

**Table 1:**
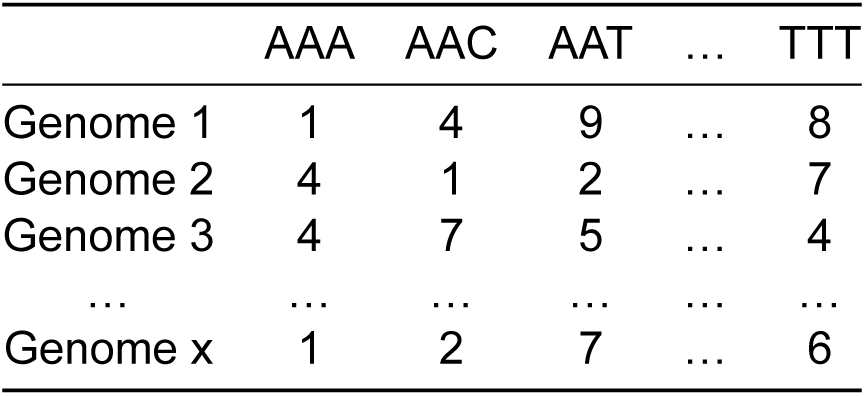
Example of a K-mer counts matrix used as input for the baseline models.

As input to the Convolutional Neural Network (CNN) and Recurrent Neural Network (RNN) models, the *k-mers-on-a-string* approach is used (fig. 1). The encoding is done in two distinctly different ways, one using one-hot embeddings of the tokens and the other using ESM-C embeddings as genome representations. For the first approach, the downsampled genomes are tokenized using fractions of the suffix length, then the resulting tokens are encoded using a one-hot encoding of the tokens (fig. 1). The ESM-C embeddings are computed by translating the entire downsampled DNA sequence into amino acids, ignoring stop-codons, then embedding the resulting protein using the ESM-C 600b embedding model (Hayes et al., 2025), resulting in a vector of size 960 per aminoacid, all of which are averaged across the rows to obtain a single vector with length 960, representing the entire genome (fig. 8). ESM-C is usually used to embed individual proteins one by one, but here we extend the method to represent the full genome sequence all at once.

### 2.4 Model architectures

Four different model architecture types were used in this analysis, namely two ensemble models and two neural network architectures. An overview of the parameters for the different architectures can be seen in table 2. The Scikit-Learn Python machine-learning module (Pedregosa et al., 2012) was used to create the machine learning architectures running the ensemble models, which use the k-mer counts and frequency matrix. The RandomForestClassifier and HistGradientBoostingClassifier modules from Scikit-Learn were used for running these two models. The HistGradientBoosting classifier is a faster Scikit Learn implementation of the traditional Gradient Boosting, inspired by LightGBM. The Python tensor module PyTorch was used to create and run the CNN and RNN models. Two different CNN models were created, one smaller and one larger, with 96 and 128 channels each and 2 and 5 hidden layers each. The small CNN had kernels of size 7 and 3, and the larger CNN had kernels of size 50, 7, and 3. The RNN was made up of a gated recurrent unit, 64 channels, and 2 hidden layers.

**Table 2:** Parameters used to build and train the different machine learning models.

|  | **RNN** | **CNN LARGE** | **CNN SMALL** | **Random Forest** | **HistGradient-Boosting** |
| --- | --- | --- | --- | --- | --- |
| **Main ML Library** | Pytorch 2.5.1 | Pytorch 2.5.1 | Pytorch 2.5.1 | Scikit-Learn 1.7.2 | Scikit-Learn 1.7.2 |
| **Channels** | 64 | 128 | 96 | NA | NA |
| **Hidden layers** | 2 | 5 | 2 | NA | NA |
| **Bidirectional** | Yes | NA | NA | NA | NA |
| **Dropout** | 0.2 | 0.2 | 0.2 | NA | NA |
| **Pooling** | Mean | NA | NA | NA | NA |
| **Optimizer** | AdamW | AdamW | AdamW | NA | NA |
| **Learning rate** | 1e-4 | 1e-4 | 1e-4 | NA | 1e-2 |
| **Weight decay** | 1e-4 | 1e-4 | 1e-4 | NA | NA |
| **L2 regularization** | NA | NA | NA | NA | 1e-3 |
| **Kernel sizes** | NA | 50, 7, 3 | 7, 3 | NA | NA |
| **Loss function** | Cross Entropy | Cross Entropy | Cross Entropy | NA | Log loss |
| **Max features** | NA | NA | NA | NA | 0.9 |
| **Python version** | 3.12.7 | 3.12.7 | 3.12.7 | 3.12.7 | 3.12.7 |
| **GPU utilization** | Yes | Yes | Yes | No | No |

### 2.5 Model training and genome partitioning

Cross-validation is used to train and evaluate the model. Before training the models, a sequence-based clustering of the genomes is performed to avoid data leakage during training and to prevent the models from memorizing the data. SourMash (Irber et al., 2024) and Scipy (Virtanen et al., 2020) are used to cluster the downsampled sequences into clusters of similar sequences. First, each token (non-overlapping k-mer) is handed to the Sourmash minhash function with the parameter n = 1000, selecting the smallest 1000 hashes as the representation for the genome. Now, Jaccard distance is calculated between all genomes using Sourmash’s implementation of the Jaccard distance function, and the results are stored in a distance matrix. SciPy’s hierarchical clustering function is used to create a clustering based on the distance matrix. The threshold is optimized to get around 30 groups. However, the number of clusters and the number of genomes in each cluster vary widely depending on sequence diversity. During model training and evaluation, the resulting clusters of similar sequences are kept together, avoiding similar sequences being split between the training, validation, and test partitions.

Cross-validation is used while training the model, after which the model is evaluated on a test set. 20 % of the clusters are selected as a test set, the remaining 80 % of the groups are used for a 5-fold cross-validation, where 60 % of the total clusters are used for training, and 20 % are used for validation in each fold. During each training fold, the validation set is used to determine when training should stop, and the test set is used to estimate how well the model generalizes to unseen data. The final model performance report consists of different performance metrics, Balanced Accuracy, and weighted F1 score averaged over the 5 folds.

### 2.6 Performance metrics

Balanced accuracy and F1 score macro are both calculated using sklearn’s implementations. Balanced accuracy is calculated by computing the weighted average recall for each class eq. (1). The F1 macro score is calculated by calculating the per-class F1 score and taking the average (eq. (2)).

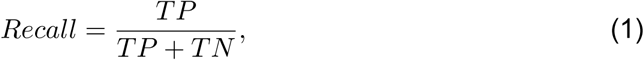

Recall score as defined in Scikit-Learn. *TP* is the number of predictions that are true positive and *TN* is the number of true negative predictions.

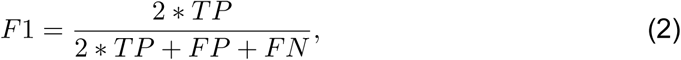

Equation for calculating the per class F1 score. *TP* is the number of predictions which are true positive, *FP* are the number of false positives and *FN* is the number of false negatives.

### 2.7 Model hyperparameters

When training the CNN and RNN models, model weights are updated continuously based on the cross-entropy loss function. Cross-entropy loss using “mean” reduction can be described using eq. (3) and eq. (4),

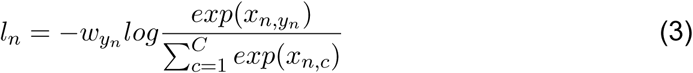

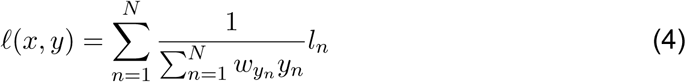

where *ℓ* is the loss, *x* is the prediction, *y* is the target label, *w* is the weight to scale the loss for, based on the number of samples in each class, and *N* is the length of the batch (Ansel et al., 2024).

The AdamW optimizer is used to optimize the model weights and biases during training based on the cross-entropy loss function. A learning rate of 1*e^−^*^4^ and weight decay of 1*e^−^*^4^ are used for both the CNN and RNN architectures (table 2).

### 2.8 Explainable features

For the ensemble models, the impact of different k-mers can be determined by computing a Shapley Analysis. This is a way to determine which features contribute most to a model’s output. The values are calculated using the SHAP Python package (Lundberg & Lee, 2017) after fitting the HistGradientBoosting classifier to the training data, and using the test data for calculating the Shapley values. The results are in the form of different charts displaying the top features (k-mers) and their impact on the model.

### 2.9 Language and Hardware

All models were run on the DTU Health Tech cluster with both CPU and GPU nodes. The CNN and RNN models were run on NVIDIA GPUs using CUDA, with between 46 GB and 96 GB of VRAM. The RandomForest classifier and HistGradientBoosting classifier only used the CPU.

Scripts for this project were written and run in Python3, with the server nodes using Python version 3.12.7. For running the ensemble models, the Scikit-Learn library version 1.7.2 was used. To run the CNN and RNN architectures, the Python tensor library Pytorch version 2.5.1 was used table 2.

## 3 Results

We will first investigate various representations of down-sampled genomes and different machine learning models to determine which down-sampling parameter combinations and genome representations yield the best results on different phenotype prediction tasks. The two datasets were chosen to get a set of different prediction tasks aiding the development of the genome representations and the machine learning models, as well as for benchmarking the models and genome representations. The *Bacformer* dataset contains a collection of bacterial genomes, retrieved from GenBank, along with their annotated phenotypic traits. The dataset was originally compiled for and used to finetune and evaluate a DNA foundation model for phenotypic traits classification (Wiatrak et al., 2025). The other dataset is a collection of *E. coli* genomes with the target of classifying whether they are resistant to the antibiotic gentamicin or not. Genomes were retrieved from the BV-BRC database (Olson et al., 2023).

### 3.1 Bacterial traits phenotyping

The *Bacformer* dataset (Wiatrak et al., 2025) consists of a range of labels with varying amounts of genomes and levels of dataset balance. We limit the analysis to the phenotypic traits (table 3), which appear in the *Bacformer* papers analysis.

**Table 3:**
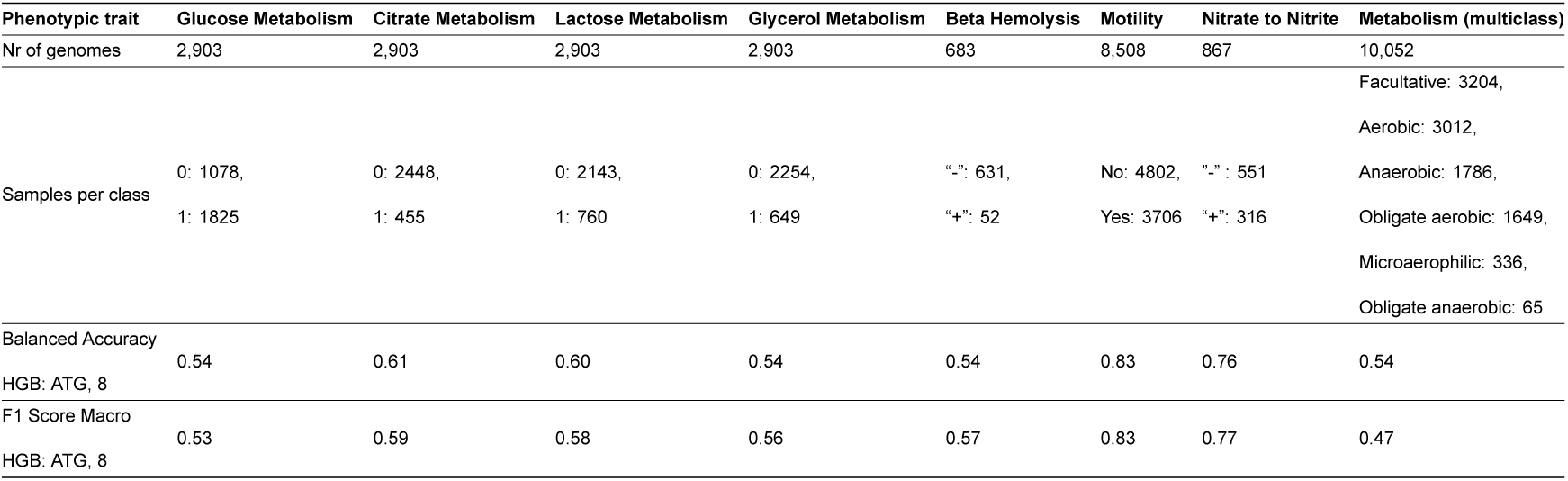
Metadata for the selection of the *Bacformer* dataset’s tasks used. Column headers describe the prediction task. The total number of genomes for each task is listed, along with the number of samples for each class. Finally, the average Balanced Accuracy and average F1 Score Macro for the HistGradientBoosting model are listed for each prediction task.

#### Finding optimal k-mer prefix and suffix length combination

To strike a good balance between the amount of genome downsampling and accuracy of the predictions, the optimal k-mer prefix and suffix length combinations were investigated. A large CNN model was trained on a task classifying whether bacteria were Gram-positives or Gram-negatives. The k-mer prefix size was varied in four steps from CGTCACA to CGTC by removing one nucleotide from the right, one by one. The suffix length was varied from 1-mers to 8-mers. The resulting heatmap (fig. 2) shows that the Balanced Accuracy (BA) strikes a good balance around k-mer prefix CGTCAC or CGTCA and k-mer suffix length 5-6. We therefore choose to primarily work with prefixes of around length 5 and suffixes of around length 6 going forward, to get good model performance without needing too much compute.

**Figure 2:**
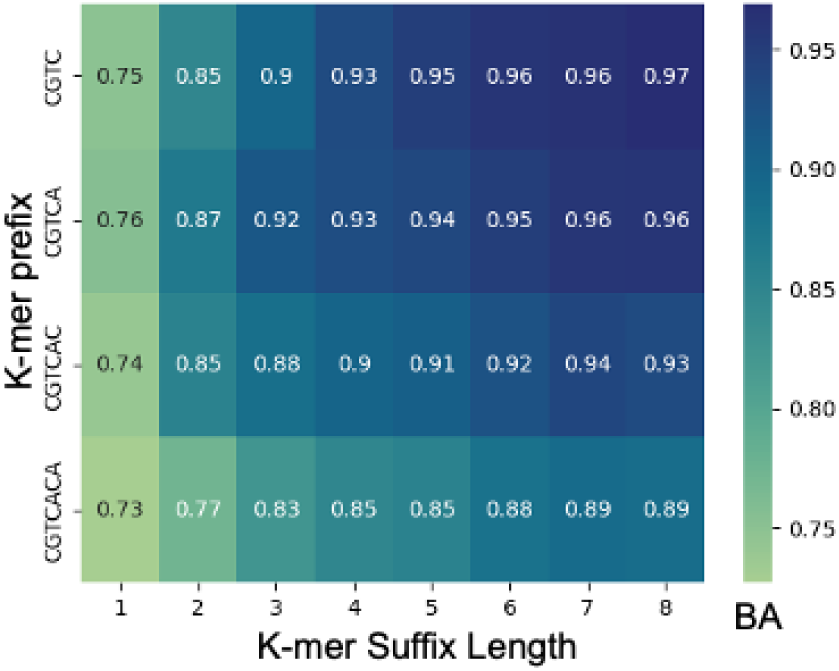
Heatmap showing Balanced Accuracy (BA) of a larger CNN model trained on the Gram staining task, comparing different combinations of k-mer prefixes, shown on the y-axis, and k-mer suffix lengths, shown on the x-axis. BA is encoded as the color of the square.

#### Tokenization strategies

Different token sizes were tested to see which tokens perform the best using both the CNN and RNN models. After downsampling the genome with prefix ACATG and suffix size of 6, each k-mer was tokenized into 1, 2, or 3 nucleotides and each token encoded as a one-hot vector (fig. 3a). This results in vectors of size 4^1^ = 4, 4^2^ = 16 and 4^3^ = 64 respectively. Performance of the different tokenizations was evaluated on the motility task. BA appears to hit a sweet spot at token size 2 for both the CNN and RNN models, but nothing definitive can be concluded, as the uncertainty is quite high and BA is calculated across only 5-folds (fig. 3b). A token size of 6 was not able to run to completion, as that representation takes up too much memory during runtime, creating a vector of size 4^6^ per 6-mer. Going forward, a token size of 1 was selected as it’s the representation with the smallest memory footprint, while still having a relatively high performance.

**Figure 3:**
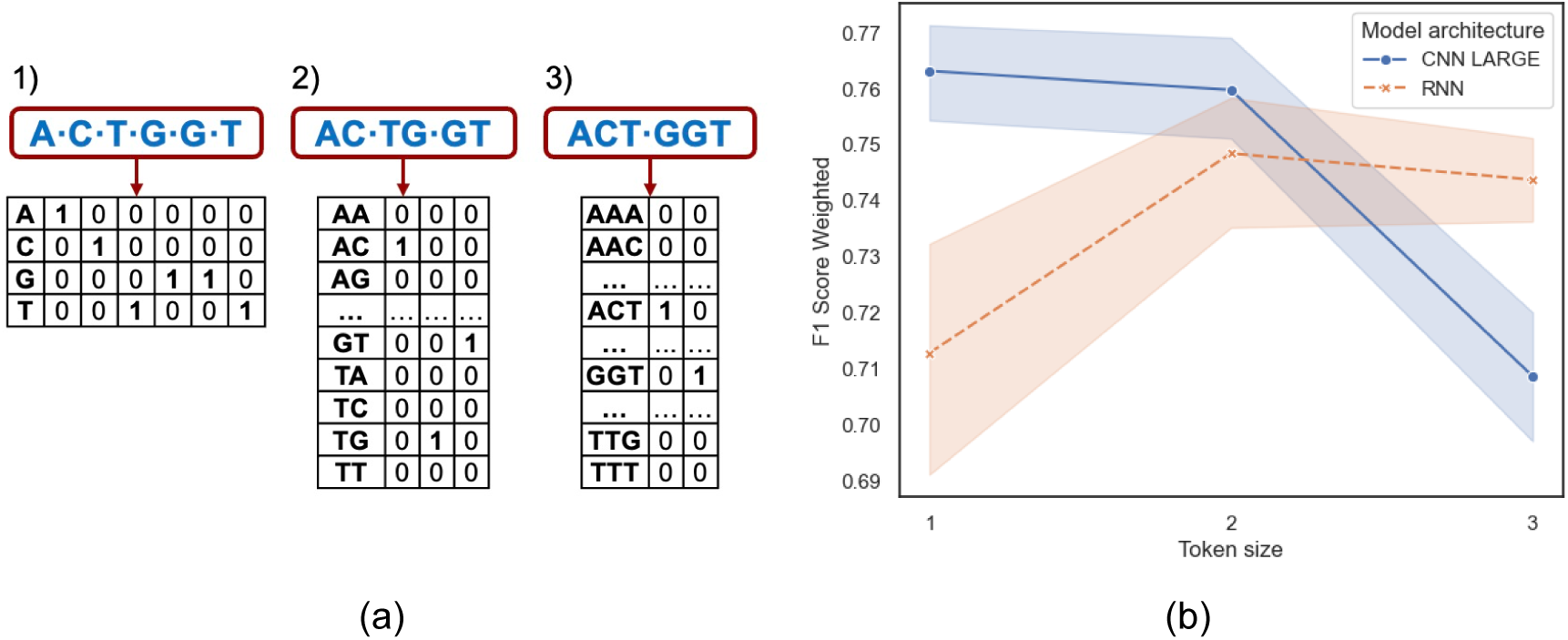
a) Illustration of the resulting input vectors of a 6-mer tokenized into tokens consisting of 1) single nucleotides, 2) 2 nucleotides, and 3) 3 nucleotides. b) Balanced Accuracy (BA) of RNN (orange cross) and CNN model (blue dot) predictions on 3 different token sizes. Models were trained on genomes downsampled using the prefix ACATG and suffix length 6 and evaluated on the motility task. Results are shown as a line plot with token size on the x-axis and BA on the y-axis.

#### Clustering downsampled genomes

We wanted to examine sequence similarity for the sequences used in the different tasks to confirm if it was necessary to create clustered partitions when training the models. For this analysis, genomes were downsampled using the prefix ACATG and a suffix length of 6 using genomes from the 4 tasks: Motility, Beta Hemolysis, Nitrate to Nitrite, and Glucose utilization. Clustering was performed by computing the distance matrix using the SourMash (Irber et al., 2024) Jaccard distance function and clustering the genomes using Scipy’s linkage function (Virtanen et al., 2020), and the average Euclidean distance. The resulting clustermaps (fig. 4) illustrate the importance of clustering the genomes before using them for machine learning model training. Especially the motility task (fig. 4a) and Nitrate conversion task (fig. 4c) clearly show clusters of similar genomes with the same class labels, indicating that clustering was indeed necessary to avoid data leakage.

**Figure 4:**
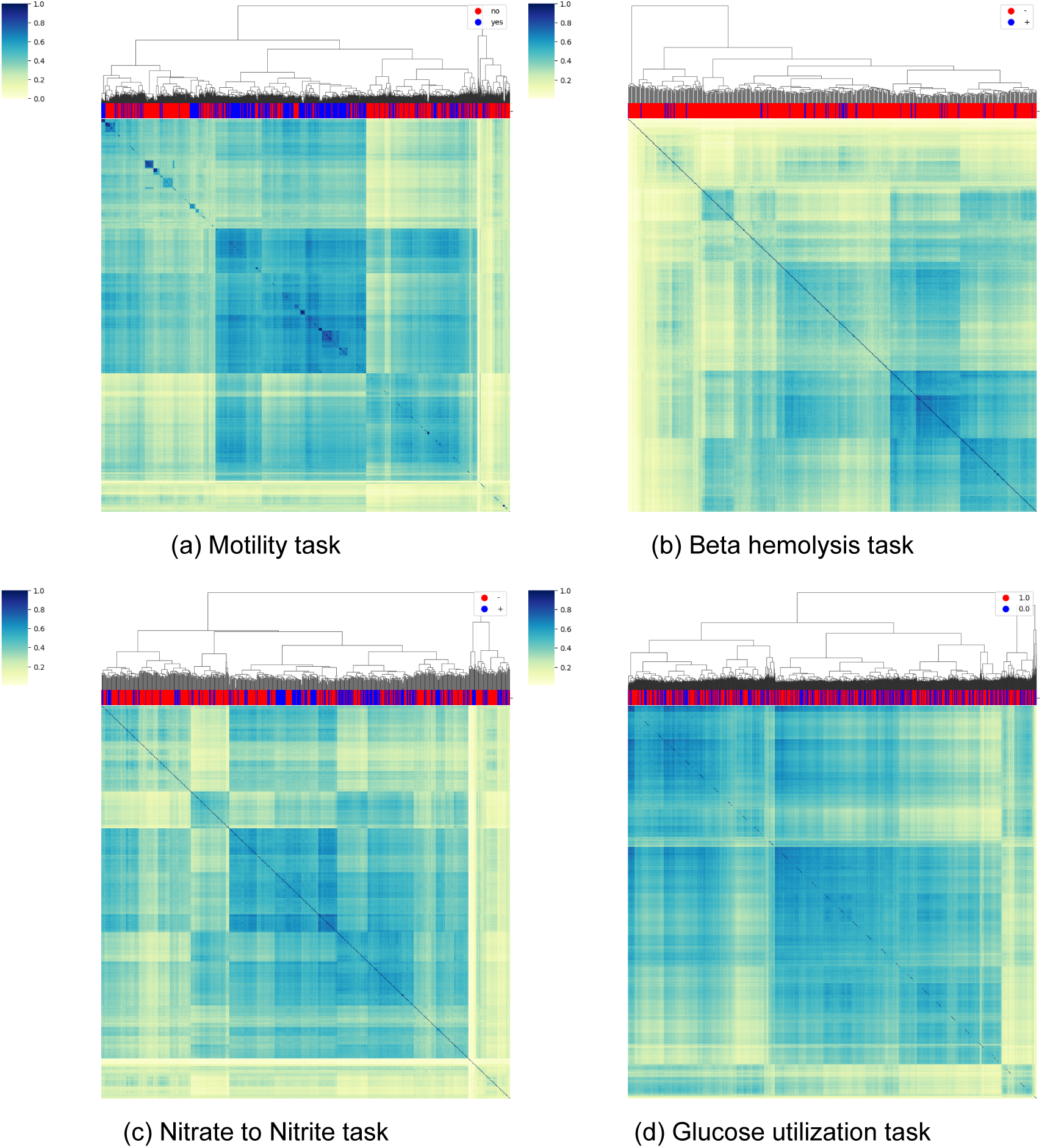
Clustermaps of the genomes from the *Bacformer* dataset (Wiatrak et al., 2025). Genomes were downsampled using prefix ACATG and suffix length 6, and distance matrices were computed using SourMash’s (Irber et al., 2024) Jaccard distance function on the 1000 k-mers with the smallest hash-values per genome. Scipy’s linkage function using the Euclidean metric and average distance method was used to create the clusters. The darker blue an area is, the closer the two genomes are, and the more light yellow the area is, the bigger the distance is between those two genomes. The red and blue bar at the top illustrates the label of the genome in that column. The linkage plot above each clustermap shows the linkage clusters as a tree.

#### Dataset partitioning methods

To compare the effect of random partitioning (Random) of the training test data versus the clustered partitioning based on genomic similarity of the genomes (Clustered), two RNN models were trained using genomes down-sampled using prefix ACATG, suffix size 6, both using the GroupKFold iterator, with one using a random partitioning and one using the clustered partitioning approach. The genomes were one-hot encoded before they were given to the model. The random partitioning method randomly distributes the training data between the training folds, not considering the distance between genomes. The clustered method keeps similar DNA sequences in one partition, meaning a group of similar sequences will be held together when creating the training, validation, and test partitions to avoid data leakage during training. No significant performance benefit was found between the random and clustered partitions (fig. 5). In the case of glycerol and citrate metabolism and nitrate reduction, along with beta hemolysis, the clustered partitions outperform the random partitions. In the other tasks, the results are the opposite. These results favor the clustered partitions, since performance is approximately identical and the clustered partitions prevent data leakage.

**Figure 5:**
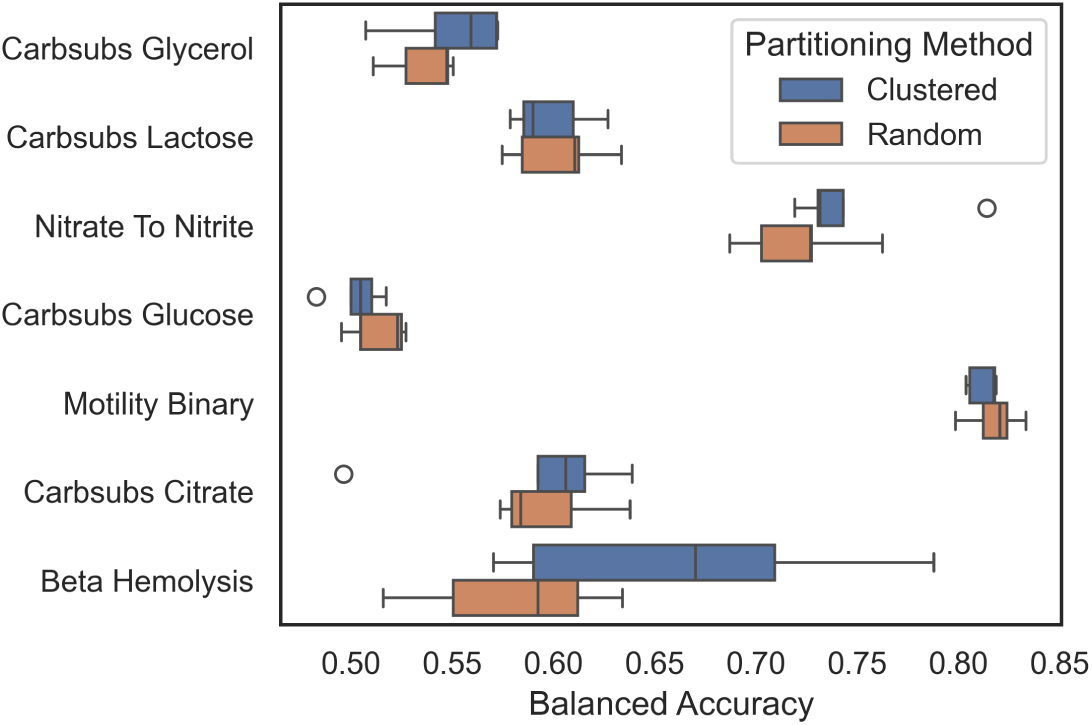
Balanced Accuracy (BA) of 7 different phenotypic traits on a clustered vs a random partitioning method. Results are from an RNN model trained across 5-folds on genomes down-sampled using the prefix “ACATG” and suffix size 6, and one-hot encoded. BA is visualized on the x-axis, with the 7 phenotypic traits listed on the y-axis. The partitioning method is encoded by color, blue for clustered and orange for random partitioning.

#### Partitioning functions

Comparing a RNN, a large CNN, and two Histogram Gradient Boosting models across two distinct training partitioning methods, GroupKFold and GroupShuffleSplit, indicates an increased BA in GroupKFold compared to the GroupShuffleSplit approach (fig. 6). This could be explained by the fact that GroupShuffleSplit overlaps the validation set across folds, which might lead to data leakage and overtraining, which the test set penalizes, giving lower overall BA scores to the GroupShuffleSplit when compared to the GroupKFold method. For these comparisons, models were trained on different combinations of prefix and suffix length. RNN and CNN models were both trained on genomes encoded using the one-hot encoding approach. The two HistGradientBoosting models were trained using frequency matrices. As GroupKFold had increased performance compared to GroupShuffleSplit, GroupKFold was selected as the data set partitioning of choice going forward.

**Figure 6:**
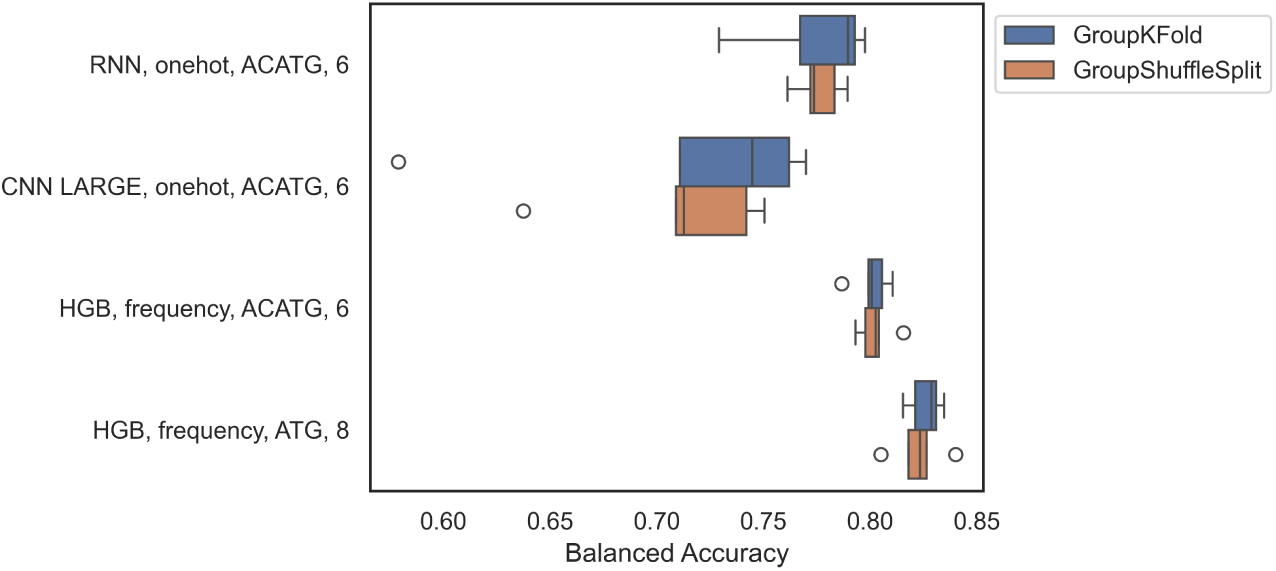
Balanced Accuracy for four different models comparing the performance of GroupKFold vs GroupShuffleSplit. The two HistGradientBoosting models were trained on downsampled genomes using frequency matrices as inputs. The RNN and CNN models were both trained on downsampled genomes using a one-hot encoding. The y-axis lists the model parameters: architecture, encoding, and the prefix and suffix lengths used for downsampling.

#### Data size impact on model performance

To investigate the impact of the dataset’s size on the different machine learning architectures’ performance, the genomes with labels for the motility task data were randomly subset from 10 % to 100 % of the genomes in steps of 10. The subset data set was used to both train and evaluate the models. The 8508 genomes in the *bacformer* dataset, which had labels for the motility task was used for this analysis. Results show that the CNN and RNN models both start with a relatively low BA using only 10 % of data and increase to BA scores of ∼0.74 and ∼0.77, respectively, (fig. 7a). The Random Forest and HistGradientBoosting models, on the other hand, both start at around BA 0.78 and only increase slightly to BA ∼0.82. This illustrates that the amount of training data impacts the RNN and CNN architectures much more than it impacts the smaller models. We speculate that with even more training data, the RNN and CNN models might reach or beat the performance of the HistGradientBoosting and Random Forest models. On the other hand, it also looks like the performance tapers off, indicating that the RNN and CNN architectures will reach a “learning upper bound” before they hit the performance of the HistGradientBoosting and Random Forest models. This should be investigated further. Analysing the average BA of the HistGradientBoosting model, trained on downsampled genomes using prefix ATG and suffix length of 8, table 3, reveals that performance does not seem to be dependent on the number of samples for a given task. Instead, it seems class balance plays a role here. Charting the BA of the HistGradientBoosting model, trained on downsampled genome representation using the prefix ATG and the suffix length 8, against the imbalance of the dataset (fig. 7b) illustrates this being the case. Generally, the more balanced the dataset is, the higher the BA is.

**Figure 7:**
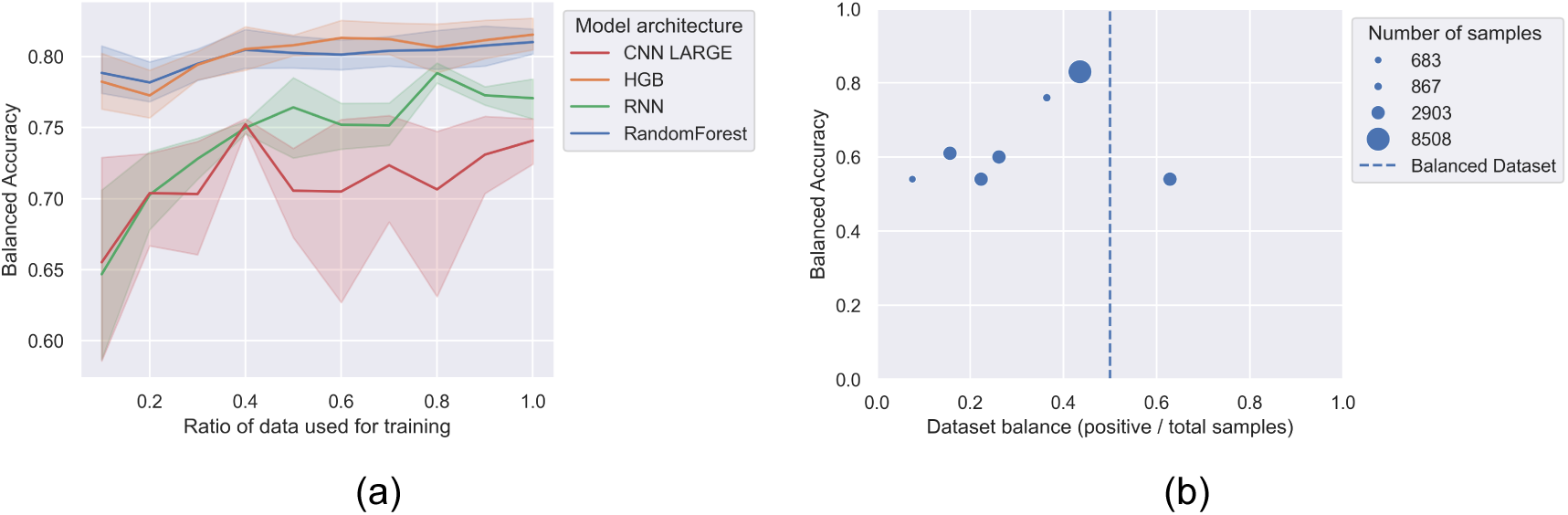
a) Impact on Balanced Accuracy (BA) depending on the number of genomes used to train and evaluate the models on the motility task. Genomes were down-sampled using the ACATG prefix and suffix length 6. BA is reported for the 5 folds, with lines showing the averages, and the shaded areas showing standard deviation. b) Impact of dataset balance on BA. Scatterplot with each point representing one phenotype prediction task from the *Bacformer* paper, with the size of each point encoding the total number of samples. The X-axis encodes the balance of the dataset, with the vertical line indicating where a completely balanced dataset would be. The y-axis displays BA of the HistogramGradientBoosting model trained on a downsampled genome representation using the prefix ATG and a suffix length of 8.

#### Direct comparison with *Bacformer* foundation model

To compare our models and genome representations against the results from the *Bacformer* foundation model, we created a similar training setup to theirs without clustering the genomes into partitions. This was done to directly compare the models and down-sampling approach used here with their results, since the *Bacformer* model (Wiatrak et al., 2025) was finetuned using a simple train, validation, test partitioning, which did not take sequence similarity into account. We expect that models trained without partitioning the dataset will perform slightly better than our previous attempts, but the drawback is that we might leak data to the model, inflating the performance metrics. To attempt to compete with the advanced transformer models from (Wiatrak et al., 2025), we used both the RNN models with genomes encoded using the one-hot approach, along with genomes embedded with the ESM-C embedding approach. For this approach, the full genome is translated to protein (including stop-codons) and afterwards embedded into a vector using the ESM-C 600b model (fig. 8) (Hayes et al., 2025).

**Figure 8:**
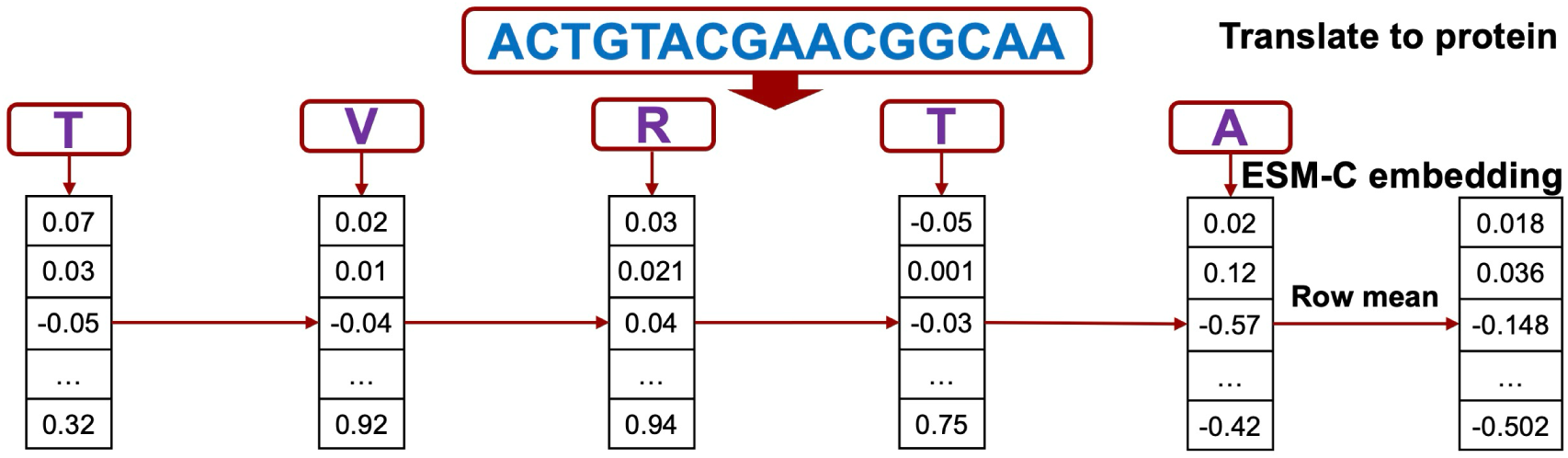
Workflow illustration of the ESM-C embedding approach. The entire DNA sequence is translated to amino acids, including stop codons, encoding each amino acid as a vector. To obtain the full genome representation, the mean of each row in the vectors is computed, resulting in one vector per genome.

When training the models using this *train, validation, test* partitioning, we see improvements to our results compared to the other methodologies tested previously. Compared to results reported by (Wiatrak et al., 2025), where the dataset was used to finetune and evaluate their model, our results look less impressive (fig. 9), as the *Bacformer* model beats the RNN models in all benchmarked tasks. However, the RNN ESM-C models get relatively close in some classification tasks, like the motility task.

**Figure 9:**
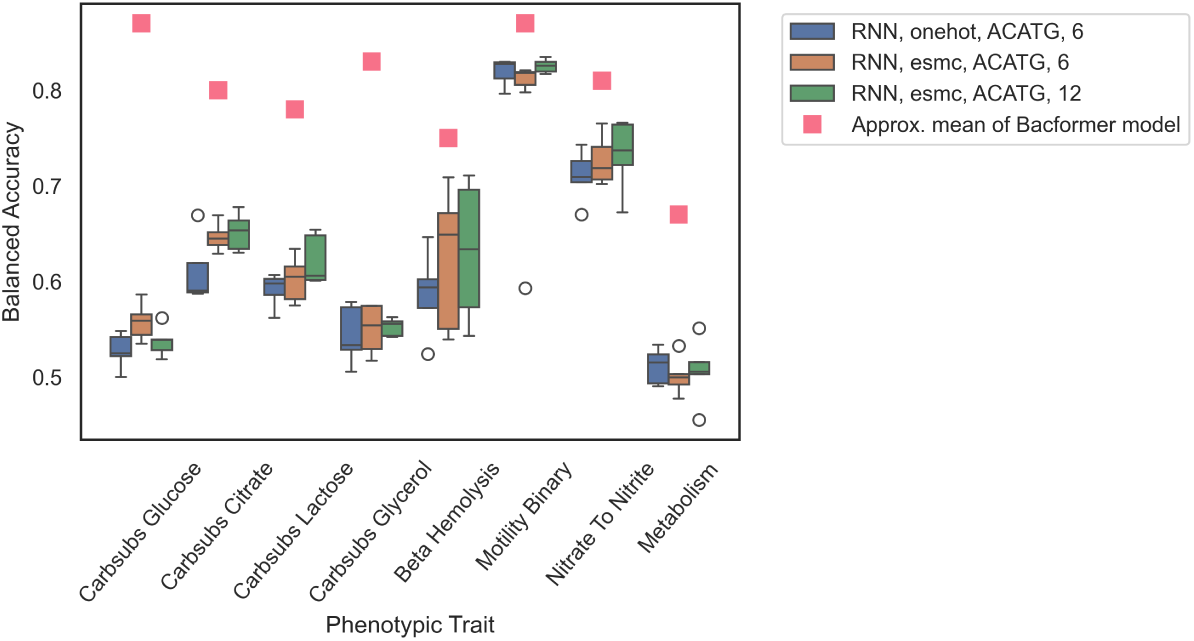
Balanced Accuracy across 8 phenotype prediction tasks over 5 validation folds for the RNN models, compared with reported results from the *Bacformer* paper, using the dataset for finetuning and evaluating a foundation model (Wiatrak et al., 2025). Three RNN models were trained on down-sampled genomes using prefix ACATG and suffix lengths of 6 and 12, respectively. The first RNN model was trained on one-hot encoded data. The other two were trained on ESM-C embeddings.

#### Model performance results

After comparing our model performance to the *Bacformer* paper (Wiatrak et al., 2025), we were interested in making a broader analysis of the performance of our different models, comparing them against each other. The models were trained on the clustered genomes using GroupKFold, and the same 60 %, 20 %, 20 % partitioning as before. The resulting BA (fig. 10) shows a performance difference between the HistGradientBoosting models and the rest of the models. In general, the HistGradientBoosting models give the best BA, especially using the prefix ATG and suffix length 8, which gets the highest BA for all tasks (fig. 10). Most models have a relatively low variance on most of the tasks. In a few of the tasks, like the beta hemolysis and motility prediction tasks, some models have higher variance. This might be explained by differences in sizes of *training*, *validation* and *test* sets and shows that some model architectures (like the CNN) are less stable on different *training* set combinations than others, with some partitions giving relatively high performance and other partitions decreasing performance drastically.

**Figure 10:**
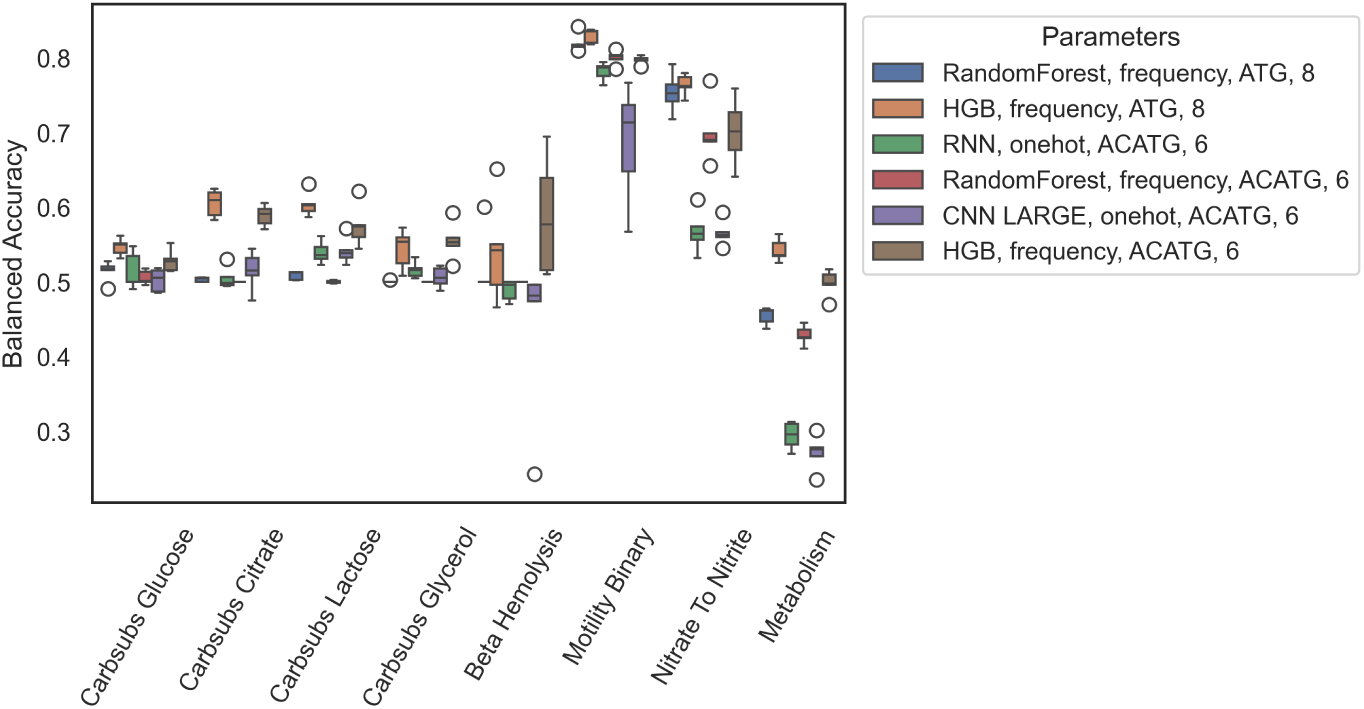
Balanced Accuracy (BA) for different models on the *Bacformer* dataset (Wiatrak et al., 2025). All models were trained using the same training, validation, and test partitions (60 %, 20 %, 20 %) using the clustered partitioning approach with the GroupK-Fold iterator. The RandomForest and HistGradientBoosting (HGB) models were trained on the down-sampled genomes using prefix ATG and suffix length of 8, along with the prefix ACATG and suffix length of 6. Finally, the RNN and CNN were also trained on downsampled genomes using the ACATG prefix and suffix length of 6. BA is reported across the 4 models and 2 different downsampling parameter combinations over the 8 prediction tasks.

#### K-mer feature extraction from HistGradientBoosting model on Motility task

When the genomes are down-sampled using the prefix-downsampling algorithm, the chosen downsampling prefix will match in many positions in the genome, resulting in a downsampled genome that will be a relatively unique representation of the entire genome. To examine which of the k-mers extracted during down-sampling had the biggest impact on model output, a SHAP analysis using the HistogramGradientBoosting model was performed.

The feature extractions obtained through the SHAP analysis on different HistGradientBoosting model shows the k-mers with the highest impact on model output. The beeswarm plot shows that these top k-mers directly impact the model output, leading the model to predict the bacteria to be either motile or non-motile depending on the k-mer frequency value (fig. 14).

In the case of the k-mer “ACATGAAAAAA”, which has the highest impact on model output in the HistGradientBoosting model, trained on down-sampled genomes using prefix ACATG and suffix length of 6 (fig. 11a), a low k-mer frequency has a negative impact on the model’s output, meaning generally that when the model sees a genome where this k-mer frequency is low, the model is directed towards predicting the 0 case, which in this case is “not motile” and when the k-mer frequency is high, the model is directed towards predicting the bacteria as “motile”. In the case of the k-mer “ACATGACTATG”, the model tend to think the genome is “not motile” when this k-mer has a high frequency according to the SHAP analysis. Analysing the contribution of individual k-mers to the model on the HistGradientBoosting model also trained on the motility task, but with genomes downsampled using parameters prefix ATG and suffix length 8, we find that some of the k-mers have a positive impact on the model when the k-mer frequency is high, directing the model towards predicting “motile”, while other k-mers have a negative impact on the model prediction, directing the model towards predicting “non motile”. If these k-mers were mapped back to the genomes, we might find that the k-mers that tend to direct the model towards predicting that a genome is motile when the k-mer frequency is high are genes coding for a protein with an impact on the bacterial motility system.

**Figure 11:**
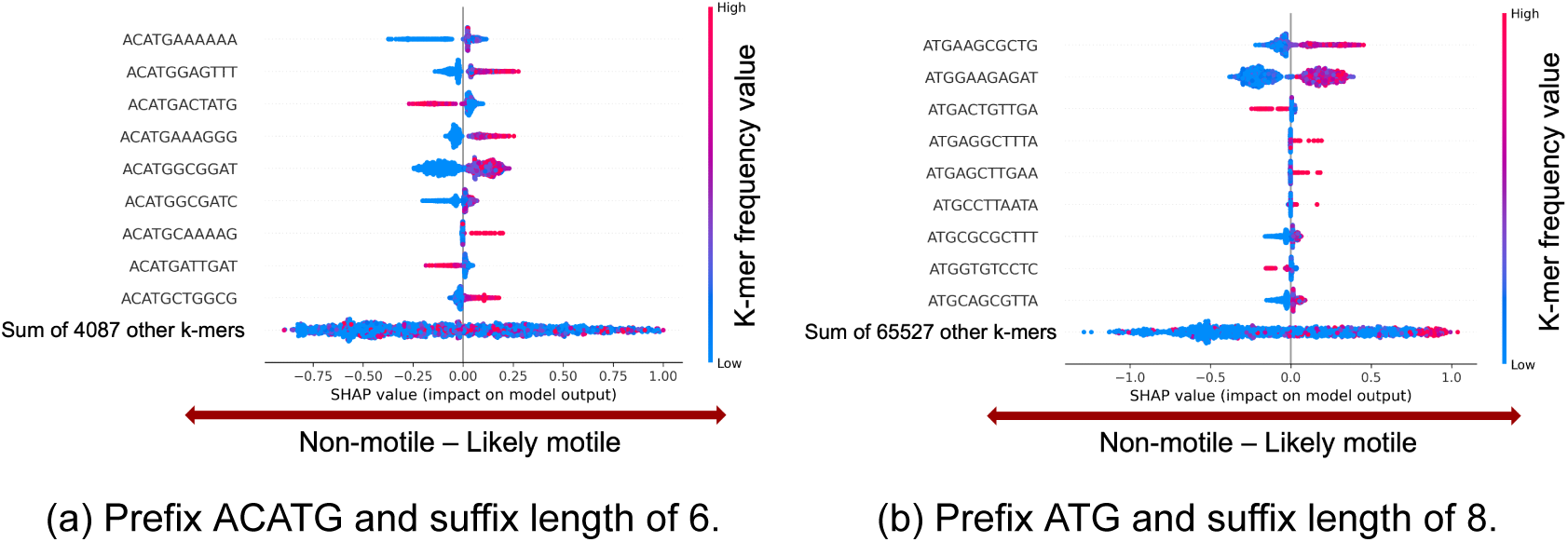
Beeswarm charts with individual SHAP values for the top 10 k-mers with the highest impact on model output for downsampled genomes using a) prefix ACATG and suffix length of 6, and b) prefix ATG and suffix length of 8, respectively. The color corresponds to the k-mer frequency value for a given genome, and the SHAP value is encoded as the position along the x-axis. In this case, a negative SHAP value corresponds to a model output likely predicting the bacteria to be non-motile, whereas a positive SHAP value will direct the model output towards predicting that the bacterium is motile.

### 3.2 Gentamicin resistance in *E. coli* and Crohn’s Disease

It has been found that the presence of adherent-invasive *E. coli*is correlated with Crohn’s disease. The gentamicin protection assay is used for quantifying invasion of epithelial cells (Kim et al., 2025), why we are interested in analysing *E. Coli* with resistance to gentamicin. We examine whether any changes can be found in the genomes of the bacteria with gentamicin resistance, apart from the gentamicin resistance gene itself, using the prefix downsampling algorithm and the machine learning models.

#### Genome clustering

While working with this dataset, we wanted to make sure that there was sufficient diversity in the genomes and that the resistance labels wouldn’t be able to simply be predicted using genomic distance. To examine this and to simply get an overview of the clusters used for partitioning the train, validation, and test sets used for training and evaluating the machine learning models, we used the same SourMash Jaccard distance matrix algorithm and clustering approach that were used to compute the clustering of the *Bacformer* dataset (fig. 4). Two clustermaps were created using different downsampling parameters. Both clustermaps (fig. 12 showed defined clusters with a high concentration of susceptible (not resistant) *E. coli* genomes, but resistance is also distributed throughout the genomes. Notably, it seems that the distance between sequences is larger in the clustermap where genomes were downsampled using prefix ACATG and suffix length of 6, (fig. 12a), since the color of the clustermap is overall more light blue than the other clustermap, (fig. 12b). As these two clustermap are made from the same original genomes, it is interesting to see that the most downsampled representation actually has a better separation of the genomes than the representation with more data.

**Figure 12:**
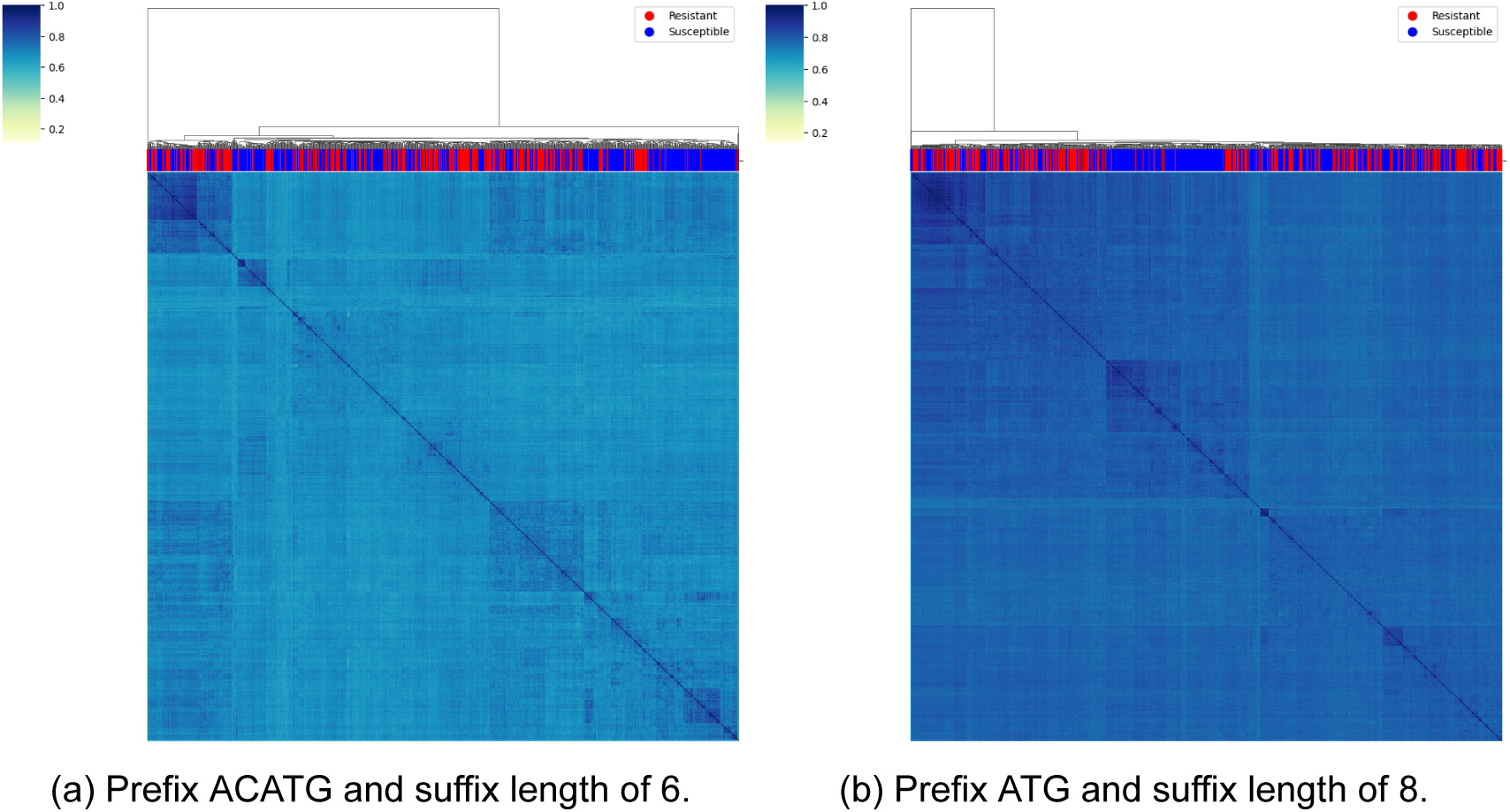
Clustermap showing clusters of the genomes from the gentamicin resistance task on downsampled genomes using parameters prefix ACATG and suffix length 6, and prefix ATG and suffix length 8, respectively. Clusters were computed using Scipy’s Linkage clustering algorithm to cluster a SourMash Distance matrix representation created using the downsampled genomes. The red and blue colored bar illustrates the class of that particular genome.

#### Overall model performance

Comparing a range of different machine learning models, genome representations, and downsampling parameters, we find that the HistogramGradientBoosting model using the genome representation created with prefix ATG and suffix length of 8 as the down-sampling parameters, by far outcompetes the other models and parameters (fig. 13). In comparison, the Random Forest model using the same exact genome representation and downsampling parameters performs much worse. The closest model performance is the HistGradientBoosting with slightly more downsampled data (prefix: ACATG, suffix length: 6) (BA=0.71) and the Large CNN (BA=0.69). The RNN model had a performance of 0.5 on this task, which means that this architecture isn’t able to learn anything about the structure of this particular data. This seems to indicate that the RNN model is not good at identifying single genes, although this has to be investigated further and might also have to do with the fact that there simply is not enough data available to train an RNN model on this task. It is interesting that the performance of the HistGradientBoosting model, trained on the downsampled genomes using prefix ATG and suffix length of 8, has the highest performance, even though the clustermaps show that the distance between genomes is smaller than the representation using parameters prefix ACATG and suffix length of 6 for downsampling the genomes (fig. 12).

**Figure 13:**
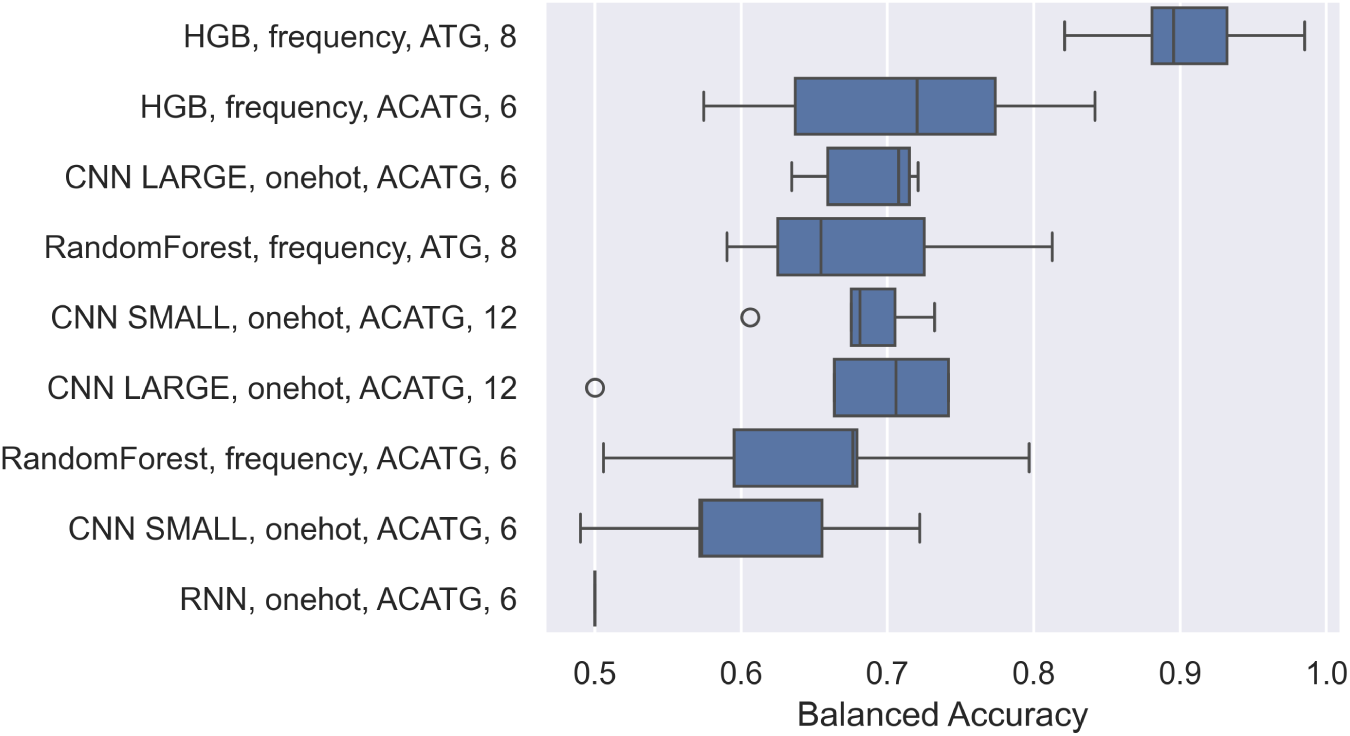
Balanced Accuracy (BA) of models trained using the 5-fold validation on different downsampled representations of the gentamicin resistance dataset. Two RandomForest models and two Histogram Gradient Boosting models were trained along with a CNN model using a smaller (2 convolution layers, 96 channels) and one using a larger size (5 convolution layers, 128 channels). Finally a RNN model with a one-hot representation was trained as well. Results for each model and parameter combination are reported as a boxplot summarizing the BA of the 5-fold cross-validation scores.

#### K-mer importance

After training the HistogramGradientBoosting baseline, we found the HistogramGradientBoosting model with genomes downsampled using prefix ATG and suffix 8 to be by far the best performing model for this prediction task (fig. 13). This result can partially be explained by a lower downsampling ratio, thereby giving the model more data, but it also seems to indicate that Antimicrobial resistance prediction tasks are very gene-specific compared to the other tasks analyzed previously, resulting in higher performance. To analyze just how gene-specific this task is, a SHAP analysis was performed on this best-performing HistogramGradientBoosting model.

Examining the mean SHAP values (fig. 14 a), we find that the 4 features (1 - 4) that contribute most to the model output are all DNA fragments mapping exactly to resistance genes, (table 4), found in the aminoglycoside database from Resfinder (Florensa et al., 2022). In comparison, none of the other top 10 k-mers matches exactly to any sequences in this database. The SHAP feature importance chart of the top 4 k-mers combined explains more of the model output than the rest of the 65533 features combined. Results shown in fig. 14 b underlines this, as the SHAP values for the top 4 k-mers from the model trained on downsampled genome representations created with prefix ATG show a clear difference between the genomes with few (or zero) of these k-mers, where those genomes likely are susceptible, and the genomes with a high frequency of these k-mers, where the genomes likely are resistant towards gentamicin. This underlines that the model learns features that can be directly traced to resistance genes, which could potentially be useful when identifying new resistance genes or possibly used to find and annotate other unknown genes in tasks where the phenotype is known but the responsible gene for the phenotype is unknown.

**Figure 14:**
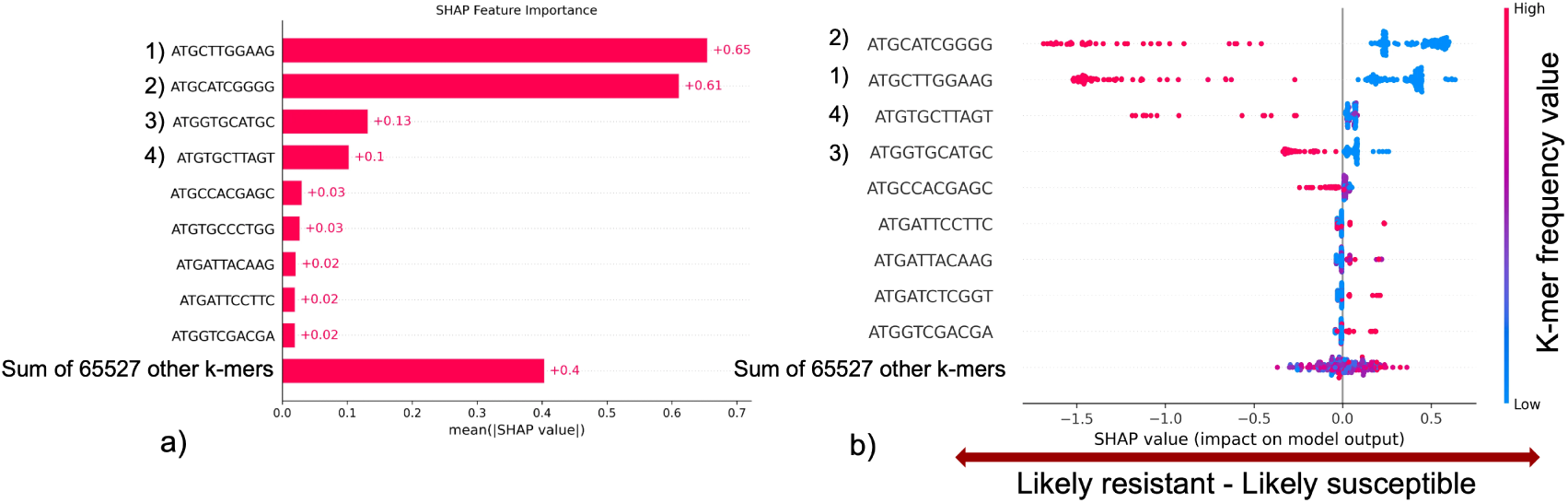
a) Bar plot showing the top 10 k-mers with the highest mean SHAP feature importance from a Histogram Gradient Boosting model being trained with prefix ATG, suffix size 8. The top 4 k-mer features match aminoglycoside resistance genes found in the ResFinder database. The rest of the most important features do not match any aminoglycoside gene in the database. This shows that the model learns features specific to resistance genes. b) Beeswarm plot illustrating the SHAP analysis from the same trained model shown in a), with each SHAP value being displayed separately as a dot. The color scale, blue to red, illustrates the impact of a specific feature being numerically low or high. A high frequency (red) of the k-mer ATGCATCGGG pushes the model’s prediction towards “resistant”. If the frequency of this k-mer is low, the model will instead tend to predict that the bacterium is susceptible to gentamicin.

**Table 4:**
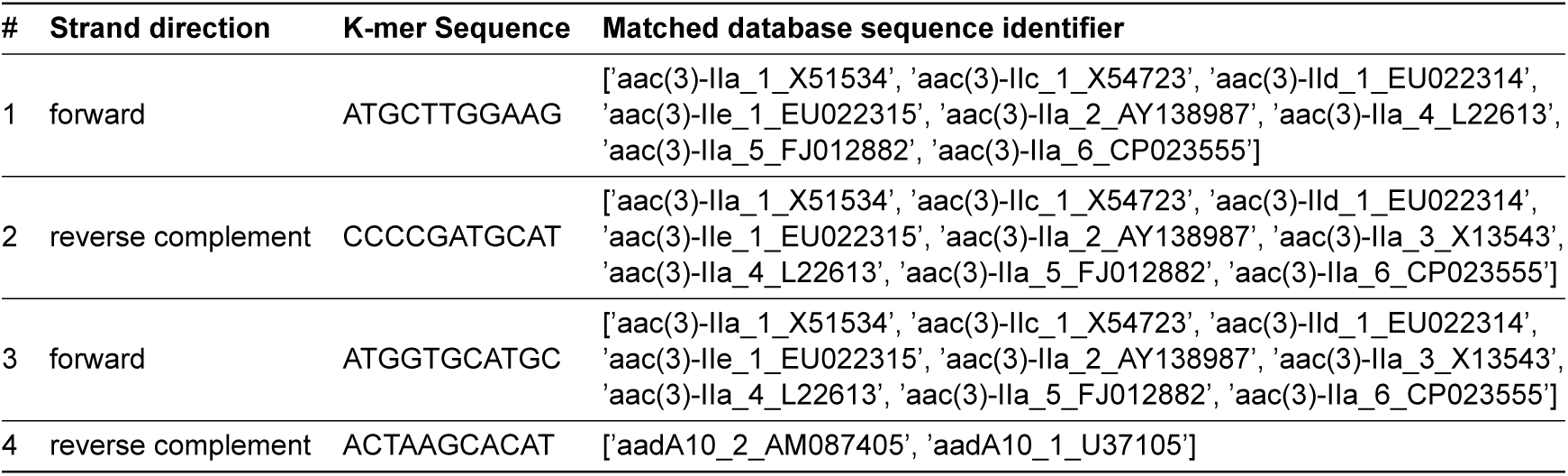
K-mer sequences obtained from the SHAP analysis, along with the identifiers of exact matching resistance genes found in the ResFinder Aminoglycoside database. Strand direction indicates whether the k-mer or its reverse complement matched the database.

## 4 Discussion

We have here demonstrated that the *prefix downsampling algorithm* can be used to downsample and represent genomes in a way that, for the most part, preserves the gene order and the structure of the genomes, creating representations that Machine Learning models can learn from. Specifically, we demonstrate that by combining the *prefix downsampling algorithm* with commonly used genome encodings, such as k-mer frequency matrices and one-hot encodings, we obtain quite powerful models and accurate predictions, particularly given the limited data and minimal computational requirements compared to more advanced foundation models. As we found, there is a price for downsampling, but in cases where using all the data is infeasible or impossible, a reduced representation may be the optimal solution to the prediction problem.

Different Machine Learning architectures and model sizes were trained and tested, but only a relatively small difference in predictive performance was found between them. This aligns with a previous study by (Kaplan et al., 2020), which demonstrates that the size and depth of a Machine Learning network have only a small effect on the loss within a wide range. They found that, up to a point, dataset size had a much greater impact on the model loss. Another study found that model improvements shifted the error, but did not shift the power-law in relation to data size, demonstrating that dataset size is more crucial than the specific model architecture (Hestness et al., 2017). This corresponds well with our findings, showing that model performance of the RNN and CNN architectures scales with the size of the dataset (fig. 7a), indicating that even more data would improve performance of these phenotype prediction models.

The HistGradientBoosting model, a derivative of the Gradient Boosting model, trained on downsampled genome representations, using the downsampling prefix ATG and a suffix length of 8, was able to predict whether *E. coli* was resistant to gentamicin with ∼ 90% Balanced Accuracy, vastly outperforming the other models and even the Random Forest model trained on the same genome representations. This demonstrates that a signal is retained after downsampling, but as we expected, the results presented also show that with more downsampling comes worse model performance. It also underlines that the choice of model architecture is very important for model performance, as the change in model architecture from Random Forest to HistogramGradientBoosting increases the performance on the same dataset drastically. As demonstrated, down-sampling can be used to find a balance between good enough predictive performance of a model and the given computational limits, be it hardware or time. Another important finding is that this model is able to learn k-mer features that can be traced directly to resistance genes (table 4). This could potentially be used to find new resistance genes or used in other places to find and annotate unknown genes when the phenotype trait is known, but the responsible gene (or genes) is not.

When comparing our downsampling algorithm and model performance with the performance of the *Bacformer* model (Wiatrak et al., 2025), none of our models were able to match the performance reported in their paper. However, it is well known that large Machine learning models are easily overtrained when the dataset is small and especially if data leakage is not controlled, as the models will simply memorize the data (Carlini et al., 2023). Since the dataset in the *Bacformer* benchmark was not carefully clustered during finetuning (to avoid very similar genomes being put both in training and in test), this might have happened here. Unfortunately, we were not able to rerun their methods with clustered training partitions, so we can only speculate, but this should be explored further, as correcting this might lead to worse performance than they report. We found that correct clustering of the data is important, as large clusters appear in many of the phenotype prediction tasks (fig. 4), which can potentially lead to data leakage if not handled properly. Additionally, we find that the clustered partitioning approach had equal or better performance compared to a random partitioning approach (fig. 5), illustrating that the clustered approach is desirable to use in any case.

Results from the models trained on the downsampled representations of the genomes from the *Bacformer* dataset predicting the 8 phenotypic traits (table 3), clearly showed the models performed best on the motility (Motility Binary) and nitrate reduction (Nitrate to Nitrite) tasks (fig. 10). This is an interesting finding as motility is governed by a very diverse set of genes and gene networks, whereas nitrate reduction is coded by genes in single operons (Rajagopala et al., 2007; González et al., 2006).

The RNN models trained on embeddings created with the ESM-C foundation model and predicting the 8 different bacterial phenotypes from the *Bacformer* benchmark (Wiatrak et al., 2025), yielded a performance improvement compared to the models trained on the same tasks using the simpler one-hot encoding. However, the embedding step was computationally very heavy and took a long time to run. Depending on circumstances, this likely is not feasible for encoding entire genomes, since it’s designed for embedding single proteins (Hayes et al., 2025). Additionally, our approach, where row means of the ESM-C embedding vectors were used to represent the entire genome, is a very crude way of representing entire genomes, which will not be able to model all genes, single-nucleotide variants, or the order of genes. We also run into another issue when using current language models for encoding downsampled genome representations, since the current models are trained on full genomes or full protein sequences, which means they may not able to explain the variance in a downsampled genome representation. This means that the ESM-C embedding model is not ideal for encoding complete genomes.

To utilize the benefits from an embedding model and tailor it for downsampled genomes, we suggest creating a foundation model trained on the downsampled genome representations, which will potentially better represent these downsampled genomes, thereby improving predictions. Such an improvement can be seen when comparing results of a transformer model trained directly on training data with a protein foundation model finetuned on the same training, which dramatically improves model accuracy (Wiatrak et al., 2025). The *Bacformer* model is an example of a Protein Language Model (PLM) that models genomes as ordered sequences of proteins (Wiatrak et al., 2025). By embedding every protein individually as a vector, this approach dramatically decreases the size of the data compared to a DNA sequence-based Genome Language Model. Additionally, since the protein language is expected to be more conserved than the DNA language, single-nucleotide variations (SNVs) in DNA will often not be detected at the protein level, potentially leading to more robust predictions, as these SNVs might act as noise for the DNA language model depending on the classification task. However, the PLM’s efficiency comes at a cost, as PLMs fail to model non-coding regions, which have been found to serve an important role in the regulation of gene expression in bacteria (Chauvier & Walter, 2024). In contrast, a Genome Language Model can capture the entire genomic context, including these regulatory elements, but requires a much larger representation than the approach used in the *Bacformer* architecture. Consequently, the optimal choice of architecture depends on factors such as whether the non-coding regions are of importance to the task at hand, as well as on the requirements for the size of the genome representation.

To improve the predictive performance of models trained on downsampled genome representations, we propose training a Transformer model on these representations using the *k-mer-on-a-string* approach. The Transformer architecture excels at learning which parts of a sequence to pay attention to, given the context around it, and can use the ordering of the k-mers for additional context and should dramatically outcompete RNNs and CNNs for this type of task (Vaswani et al., 2023). Additionally, we propose investigating the Mamba and Hyena DNA architectures, which are based on State Space Models, a generalization of the RNN architecture. Our approach may be an alternative to using the Mamba architecture, or it might be that our downsampling approach could be combined with the Mamba architecture to get an even more performant model. These architectures can create Large Language Models with a similar level of performance to the Transformer architecture (Dao & Gu, 2024), but with a much larger context window, which can be used to keep most of, or in many cases, entire bacterial genomes in memory when training. Today, most methods using Transformers are protein language models (pLMs), which involve translating individual genes from the genome to protein and embedding each protein (Wang et al., 2025). This is an effective method for phenotype prediction, but it might still discard relevant context for the models, like non-coding regions and SNVs. Only recently have genome language models (gLMs) emerged, like the Nucleotide Transformer architecture (Dalla-Torre et al., 2025), but even these are also usually only trained on parts of DNA sequences and not on the entire genome.

### Effectiveness on different phenotype prediction tasks

We believe that, depending on the phenotype prediction task, different genome representations are optimal. In the case of phenotypes resulting from gene single-nucleotide variants (SNVs), the prefix downsampling representation is likely not sensitive enough, as only a small subset of these SNVs will be detected. On the contrary, gene order is relatively easy to detect, as the ordering is kept during downsampling and is kept in the *k-mers-on-a-string* models, which the CNN and especially RNN architectures should be able to utilize. Similarly, gene copy number and the presence or absence of a gene can be estimated from the downsampled representation, even using the k-mer frequency matrices, as the frequency of a specific k-mer naturally will be correlated with the gene copy number. Our results show that the k-mer frequency matrix work well for most phenotype prediction tasks analysed.

Some bacterial phenotypes occur due to a stochastic phenotype switch or when the bacteria detect an environmental change and respond to it by switching to a more favorable phenotype (Byrne et al., 2025). These subpopulation phenotypes might not improve the fitness of those bacteria at that moment, but benefit the bacterial population if a sudden shift in environment occurs, as some of these subpopulations might thrive in the new environment. Gene switching might not be easily predictable at a gene level, since these phenotypes can be regulated by a cascade of transcriptional “switches”. Examples of this are seen in *Salmonella Typhimurium*, where the flagellar genes, responsible for motility, are regulated through a cascade of three switches (Byrne et al., 2025) and will most likely require transcriptional data to be predicted.

A different approach would be to use a “word-based” or “sub-word-based” tokenizer like the Byte-Pair Encoder (BPE). A BPE can be trained to recognize pieces of DNA that often occur together across many genomes (Sanabria et al., 2024). This would result in an encoding of each genome containing long pieces of DNA that occur often, and shorter pieces of DNA that only seldom occur in genomes, creating a smaller genome representation while keeping all the genomic information.

MinHashing and FracMinHashing algorithms are used in bioinformatics today to store DNA and RNA sequences as fractional representations, which can be used to identify genomic data sets with common sequences and to estimate genomic distance (Irber et al., 2024). The prefix downsampling algorithm can be used in a similar way to create downsampled genome representations for estimating genome distance, clustering, etc. Prefix filtering is similar to FracMinHash approach in that the downsampled genome representations naturally scale with the size of the original genome. This scaling property can be useful, and the SourMash Python package actually has an option for scaling the number of k-mers with genome size (Irber et al., 2024) which for instance is used in metagenomics. One may also envision a strategy (that may be coined SmallHash) that like the prefix filtering is able to determine if a K-mer is part of the final set on the fly by checking if it is smaller than a given threshold, for example one defining the lowest percent of the posible hash values. Fraction based sampeling may be a disadvantage for some use-cases, as it is not possible to exactly control the size of the resulting k-mer matrix, which the traditional MinHashing approach allows.

## 5 Conclusion

We have demonstrated that when genomic data is downsampled strategically through prefix downsampling, it enables the creation of efficient models that reduce the computational requirements of phenotype prediction models along with current Genome Language Model (GLM) and Protein Language Model (PLM) approaches.

By drawing inspiration from the prefix downsampling strategy of (Larsen et al., 2014) and the information about k-mer size efficacy shown by (Aytan-Aktug et al., 2020), we developed a novel downsampling algorithm. To the best of our knowledge, this represents the first application of a prefix-based or hash based downsampling of entire genomes for use in Machine Learning prediction tasks.

Our analysis of this novel approach yielded two primary insights regarding architecture and data representation. Firstly, we confirmed that ensemble models such as Random Forest and Gradient Boosting are exceptionally effective at utilizing k-mer frequency matrices for predicting phenotypic traits. Consistent with literature on tabular data (Grinsztajn et al., 2022; Xu et al., 2021), these ensemble models outperform more complex models, especially when data is limited and when genomes are very similar. Ensemble models have much fewer parameters to tune, making it simple and fast to get good prediction results, showing once again the power of simpler models (Xu et al., 2021).

Secondly, while k-mer frequency matrices are efficient for use with ensemble models, we argue that the *k-mers-on-a-string* approach, combined with advanced sequence-dependent architectures, enough data, and an efficient downsampling strategy, will unlock higher-performing models going forward. While we primarily focused on the down-sampling algorithm itself and using the downsampled genomes as input to Machine Learning models, we believe that this data structure has many other applications, as it can be used as an alternative to current MinHashing algorithms and as input for small Genome Language Models using either Transformer or Mamba architectures.

In summary, this thesis provides a framework for downsampling large genomes while retaining genomic information and discarding redundancy. By showing that a down-sampled genome representation can yield high-performance predictive models, we suggest a way toward small Genome Language Models that will enable the processing of much larger genomic databases and enable this analysis on standard computing hardware.

## 6 Code Availability

Code developed for this thesis is available at https://github.com/TorbjornBak/ML-bacterial-phenotyping.

## Notes

### Competing Interest Statement

The authors have declared no competing interest.

### Funding

- Novo Nordisk Foundation: NNF23SA0087056

